# Study of the molecular nature of resistance to bifenazate in a *Tetranychus urticae* Koch Laboratory Strain

**DOI:** 10.64898/2026.03.18.712698

**Authors:** Elena Sergeevna Okulova, Dmitrij Dmitrievich Skrypka, Olesja Denisovna Bogomaz, Roman Romanovich Zhidkin, Galina Petrovna Ivanova, Irina Anatolyevna Tulaeva, Xingfu Jiang, Tatiana Valeryevna Matveeva

**Author notes:** Corresponding author: Tatiana Valeryevna Matveeva, Full institutional address: Department of Ecological Genetics, Center for Biological Regulation of Pesticide Use, All-Russian Institute of Plant Protection, 196608 St. Petersburg, Russia.

## Abstract

**BACKGROUND:** The two-spotted spider mite, *Tetranychus urticae* Koch, is a major agricultural pest with a rapid propensity for developing acaricide resistance. Bifenazate targets mitochondrial cytochrome b (CYTB). While the G126S mutation is frequently associated with resistance, its independent role remains unclear as it often occurs with other substitutions. This study explores the molecular basis of bifenazate resistance in a Russian laboratory strain derived from a St. Petersburg greenhouse population.

**RESULTS:** Disruptive selection with increasing bifenazate concentrations generated resistant and susceptible isofemale lines. AlphaFold2 structural modeling of CYTB indicated that G126S causes a steric clash, leading to conformational destabilization, whereas other reported mutations primarily affect the ligand-binding pocket. Oxford Nanopore sequencing revealed a very low initial frequency of the G126S allele (<1%; 226/35,895 reads) in the unselected population. After one year of stepwise selection (0.00005–0.031% a.i.), the mutant allele frequency surged to 90% (7,272/8,056 reads). No other known resistance-associated mutations were found in the analyzed *cytb* fragment.

**CONCLUSION:** We report the first identification of the G126S mutation in a Russian *T. urticae* population and demonstrate its rapid fixation under bifenazate selection. Within this genetic background, G126S alone appears sufficient to confer high-level resistance, emphasizing the population-specific nature of resistance evolution and the critical need for local monitoring.

## 1 Introduction

The two-spotted spider mite (TSSM) *Tetranychus urticae* Koch (Acari: Tetranychidae) is one of the most destructive herbivorous pests; it damages more than 1,100 plant species, about 150 of them are important crops, including vegetables, fruits, and ornamentals.^1^ It has a short life cycle, high offspring production and a remarkable ability to develop pesticide resistance.^2–4^ *T. urticae* females may produce over 100 eggs of different size depending on the embryo sex: males develop from unfertilized eggs while females derived from fertilized ones.^5^ All of the above shows the relevance of research aimed at improving methods for controlling the populations of these pests.

Spider mites are most effectively managed when multiple control strategies (mechanical, cultivational, biological, and chemical controls) are used to reduce pest populations and maintain plant health.^6^ When all other control methods have failed to keep the populations under control, chemical acaricides may be needed. Chemical control of TSSM has been only partially successful, as TSSM can rapidly develop resistance to most types of insecticides and acaricides because of its short life-cycle, resulting in rapid population growth.^3^ Acaricide resistance can develop through multiple mechanisms, including enhanced metabolic breakdown of acaricides, target site insensitivity, and behavioral resistance.^7^ One of the most important mechanisms for acaricide resistance development in *T. urticae* is target site insensitivity, where single nucleotide polymorphisms (SNPs) can result in an alteration of an amino acid sequence and the translated protein which, in-turn, binds the insecticide weaklier or not at all.^8^

*T. urticae* has been documented to have developed resistance to over 96 acaricidal/insecticidal active ingredients.^9^ Tolerance to acaricides can result after a few applications.^10^ Furthermore, *T. urticae* can become fully resistant to new acaricides within two to four years, meaning that control of multi-acaricide resistant *T. urticae* has become increasingly difficult.^11^

Bifenazate (D2341, N′-(4-methoxy-biphenyl-3-yl) hydrazine carboxylic acid isopropyl ester) is a hydrazine carbazate acaricide that was discovered in 1990 by Uniroyal Chemical. It is used to control all growth stages of *Tetranychus* spp. on a wide variety of crops.^12^ Bifenazate was shown to be converted by carboxyl/cholinesterases into diazene and an activation pathway has been suggested.^13,14^

Bifenazate target is mitochondrial electron transport (MET) chain.^3,15^ The MET chain consists of four large transmembrane enzyme complexes (Complexes I–IV) that work together to transfer electrons from NADH and succinate to molecular oxygen.^15,16^ Cytochrome c oxidoreductase or the bc1 complex, catalyzes the electron transfer from reduced ubiquinone to cytochrome c.^16^ Complex III consists of three core subunits: cytochrome b (*cytb*), cytochrome c1, and the Rieske iron-sulfur protein.^15,17^ Notably, CYTB is the only complex III subunit that is encoded by mitochondrial DNA. CYTb contains two distinct quinone-binding sites, Qi and Qo, which are located near the inner and outer sides of the inner mitochondrial membrane, respectively.^18,19^

Furthermore, a strong correlation was found between specific *cytb* point mutations and the occurrence of resistance (Table 1).^12^

**Table 1.**
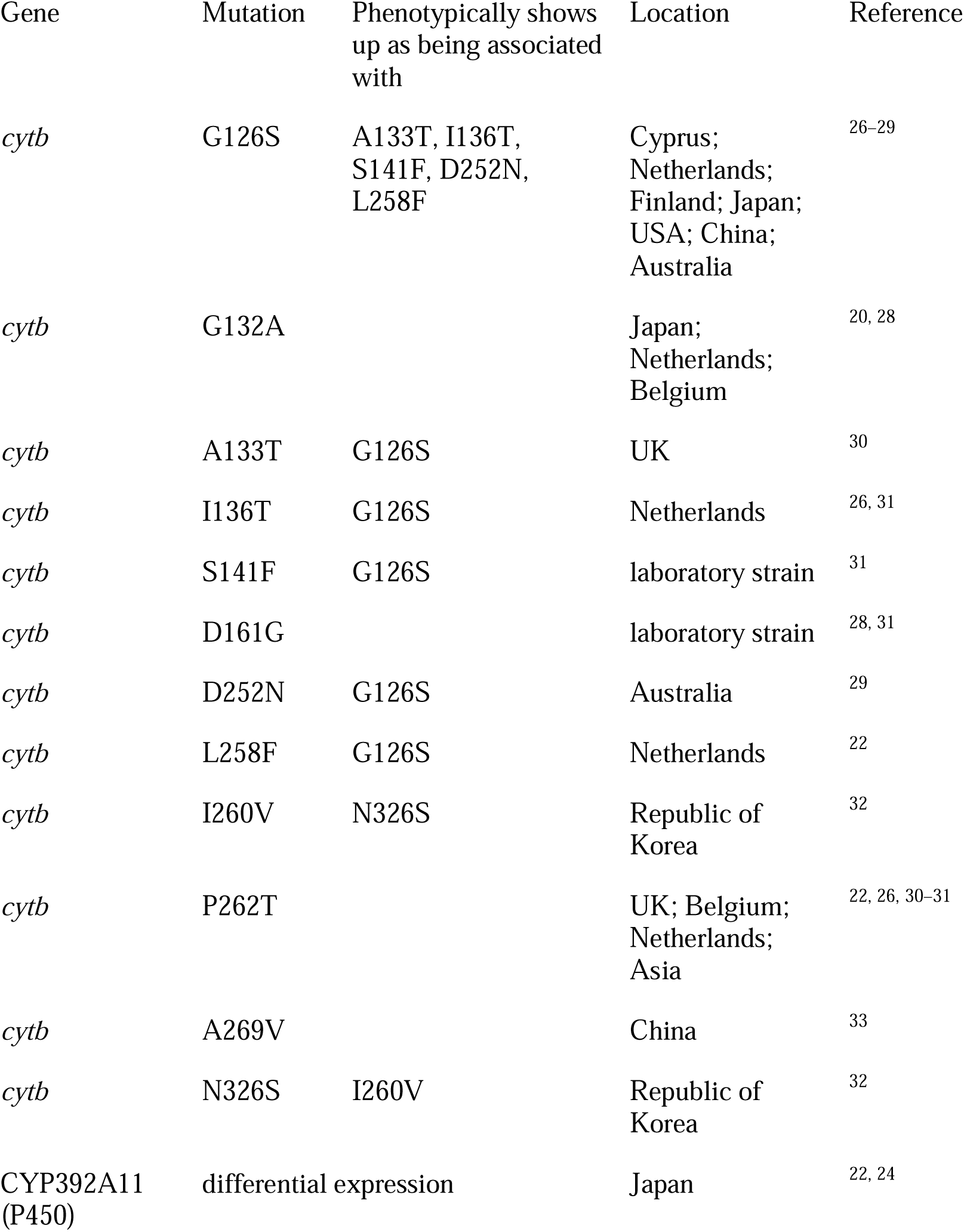
Mutations in the two-spotted spider mite *cytb* gene, associated with bifenazate resistance.

| Gene | Mutation | Phenotypically shows up as being associated with | Location | Reference |
| --- | --- | --- | --- | --- |
| *cytb* | G126S | A133T, I136T, S141F, D252N, L258F | Cyprus; Netherlands; Finland; Japan; USA; China; Australia | 26–29 |
| *cytb* | G132A |  | Japan; Netherlands; Belgium | 20, 28 |
| *cytb* | A133T | G126S | UK | 30 |
| *cytb* | I136T | G126S | Netherlands | 26, 31 |
| *cytb* | S141F | G126S | laboratory strain | 31 |
| *cytb* | D161G |  | laboratory strain | 28, 31 |
| *cytb* | D252N | G126S | Australia | 29 |
| *cytb* | L258F | G126S | Netherlands | 22 |
| *cytb* | I260V | N326S | Republic of Korea | 32 |
| *cytb* | P262T |  | UK; Belgium; Netherlands; Asia | 22, 26, 30–31 |
| *cytb* | A269V |  | China | 33 |
| *cytb* | N326S | I260V | Republic of Korea | 32 |
| CYP392A11 (P450) | differential expression |  | Japan | 22, 24 |

Due to the characteristics of mitochondrial inheritance—namely, the absence of recombination and exclusive maternal transmission—new mutations in mitochondrial DNA are rapidly fixed in populations through a bottleneck effect.^20^ This process accelerates the spread of resistance alleles, especially under strong selective pressure caused by the widespread use of bifenazate.

Metabolic resistance is generally associated with enhanced activity of enzymes that detoxify acaricides before they reach their target sites. This mechanism is driven by differential expression of detoxification genes, which can result from changes in regulatory elements (cis- or trans-regulation) or gene duplication.^21^

Cytochrome P450 enzymes (CYPs) are the most extensively studied detoxification enzymes in various species, including insects and mites. Most functionally validated CYPs belong to the CYP392 family, which exhibits increased diversity in *T. urticae.*^21^ For instance, the P450 enzyme CYP392A11 is capable of metabolizing bifenazate through hydroxylation of its ring structure.^22^ This enzymatic modification significantly reduces the toxicity of bifenazate, thereby contributing to the development of resistance.

Despite frequent increases in the expression of detoxification genes in response to pesticides, the molecular mechanisms underlying these changes remain largely unclear. Researchers rarely succeed in identifying mutations that directly trigger the upregulation of these enzymes.^20^

The high levels of resistance observed in some populations of *T. urticae* are likely the result of a combination of mechanisms, including target-site mutations and enhanced detoxification mediated by P450 enzymes.^22–25^ These mechanisms may act synergistically, significantly increasing resistance levels compared to the effects of each mechanism individually.^22^

SNPs frequently interact with other genetic and physiological factors. This complicates the study of their role in isolation and the interpretation of research results. Of particular interest is the G126S mutation in mitochondrial cytochrome b, which has long been considered an important factor responsible for resistance in *T. urticae* to bifenazate and related compounds. However, later studies based on sequencing and functional tests indicate that this mutation is unlikely to be the sole cause of resistance, and its effect appears only in combination with other mutations.

For example, one study showed that lines of the mite *Panonychus citri* with the G126S mutation exhibit only moderate levels of resistance to bifenazate, whereas the combination of G126S with an additional A133T mutation significantly increases resistance.^34^ In another study, cross-resistance in a *T. urticae* line carrying the G126S + S141F combination conferred high levels of resistance to both acequinocyl and bifenazate. In contrast, a line with the single G126S mutation remained susceptible to acequinocyl, casting doubt on its role in resistance mechanisms.^35^

Additional data showed that G126S was also detected in *T. urticae* populations susceptible to bifenazate. This led to the hypothesis that this mutation is not a reliable marker for bifenazate resistance. Nevertheless, resistance was observed in populations where G126S occurred in combination with I136T and S141F mutations. It was also noted that five mutations conferring bifenazate resistance (A133T, I136T, S141F, L258F, D252N) were always found alongside G126S, yet none of them appeared independently or in other combinations.^29^

The complexity of interpreting the role of G126S is further compounded by its interaction with metabolic detoxification mechanisms. Experiments selecting acequinocyl-resistant populations revealed the G126S mutation; however, it was also detected in bifenazate-susceptible populations, calling into question its key role in resistance to this compound. While resistance to acequinocyl showed maternal inheritance, resistance to bifenazate depended on additional factors. Studies using synergists, such as piperonyl butoxide (PBO), identified the role of cytochrome P450 enzymes in enhancing resistance, supporting its polygenic nature and the interaction of mutations with metabolic mechanisms.^24^ Finally, it was concluded, that the G126S mutation may serve as a marker of resistance, but its effect largely depends on the presence of other mutations or metabolic detoxification mechanisms. These findings emphasize the need for a comprehensive approach to studying resistance, incorporating analyses of both genetic and physiological factors and their interactions.

To our knowledge, no studies have been conducted on the molecular genetic control of resistance of Russian spider mite wild populations and/or laboratory cultures to bifenazate. In the framework of this study we have investigated the molecular mechanisms of bifenazate resistance in laboratory cultures of TSSM from All-Russian Institute of Plant Protection, initiated from mites, collected in greenhouse of JSC “Flowers” in St. Petersburg from a Solo rose.

## 2 Materials and methods

### 2.1 TSSM cultures

Experiments were conducted on females of the common spider mite *Tetranychus urticae* Koch. Mite cultures were selected using a diagnostic concentration of the acaricide bifenazate, a carbazate. The line was developed from a laboratory culture selected with dimethoate (BI-58 40 % e.c.), originally brought from the greenhouse of JSC “Flowers” in St. Petersburg from a Solo rose.

### 2.2 Maintaining of TSSM

Homozygous laboratory strains of *T. urticae* were obtained by disruptive selection during inbred propagation. Resistant (R) and susceptible (S) strains of mites were cultured as separate colonies on isolated leaves of the common bean (Phaseolus vulgaris) placed in a cuvette on a layer of moistened cotton wool. The mites were maintained at 22°C and 18 hours of daylight in controlled environments designed to optimally support their development.

Mites were treated by immersing infested leaves in an aqueous solution of the toxicant bifenazate, specifically using the commercial pesticide Fenobi at a concentration of 240 g/L SC. The treatment concentration was maintained at 0.00005 %, resulting in a 75 % mortality rate among female mites. Mite colonies exhibiting 90–100 % survival after acaricide treatment were used to create R-lines, while S-lines were developed through sister selection from colonies that experienced 100 % mortality following treatment. The offspring of a single female, demonstrating either resistance or sensitivity, were individually placed on leaf rafts after bifenazate treatment to obtain age-matched offspring. The PCR experiment involved females treated with bifenazate from 1–3 rafts, showing the lowest mortality rate for R-strains and 90–100 % for S-strains. Adult females that had undergone two molting stages (proto- and deutonymph) were used in the study, which was conducted every 15–20 days, depending on the season.

### 2.3 Selective treatments

Treatments were conducted on whole bean plants grown in special containers filled with a manganese solution. When the plants began to wilt or became heavily infested with mites, a new container with freshly sprouted plants was placed nearby to encourage the pests to migrate to the new shoots. The treatments involved spraying the infested plants, with the frequency varying from 8 to 15 days, depending on how quickly the mite population recovered to the level necessary for the next treatment. Every eighth treatment, the concentration of the product was increased fivefold. In total, 28 treatments were administered, with the bifenazate concentration rising from 0.00005 % to 0.031 % of the active ingredient over the entire period.

### 2.4 DNA isolation and PCR

An adult female mite was transferred to a separate 0.2 ml tube containing 20 μl of water and heated to 90°C for 10 minutes. The solution was then carefully pipetted and either used immediately as a PCR template or stored at −20°C.

For PCR, a reaction mixture was prepared using Taq polymerase. Each 20 μl sample contained 8 μl of water, 10 μl of 2x Dream Taq Green PCR Master Mix (Thermo Scientific, Lithuania), 5 picomoles of each forward and reverse primer (BifUnM-F: 5′-AATTATAGGATCCGCTTTTATTGGG-3′; BifUnM-R: 5′-TGGTACAGATCGTAAAATTGCGT-3′), and 1 μl of template. PCR was performed in a GeneExplorer Thermal Cycler 96 x 0.2 ml (Bioer, China). The amplification program included a 3-minute preincubation at 95°C, followed by 40 cycles of 15 seconds at 94°C, 30 seconds at 60°C, and 1 minute at 72°C.

The PCR products were separated electrophoretically in an agarose gel, and the results were visualized using the GelDoc Go gel documentation system (Bio-Rad, USA).

### 2.5 Real-time PCR

For real-time PCR, a reaction mixture was prepared using the SYBR®Green intercalating dye. The 20 µl reaction mixture consisted of 8 µl of water, 10 µl of SsoAdvanced Universal SYBR®Green Supermix (Bio-Rad, USA), 5 picomoles of each forward and reverse primer, and 1 µl of template. To detect the G126S mutation, we employed the diagnostic system described by Maeoka et al.^36^, which includes two pairs of primers: allele-specific and universal. RT-PCR was conducted using a Gentier 96R Real-Time PCR System (Tianlong, China). The amplification program comprised a 3-minute preincubation step at 95°C, followed by 40 cycles of 15 seconds at 94°C, 30 seconds at 61°C, and 40 seconds at 72°C.

### 2.6 Oxford nanopore sequencing and data analysis

A MinION Mk1B nanopore sequencer (Oxford Nanopore Technologies) was employed for sequencing the amplicons, utilizing a R10.4.1 flow cell. Libraries were prepared using the Native Barcoding Kit 24 V14 (SQK-NBD114.24), and sequencing was conducted over a period of 120 hours.

Primary data processing was performed using Guppy version 6.5.7 (Oxford Nanopore Technologies). The superprecision model (SUP) was applied to convert electrical signals into nucleotide sequences (basecalling), and samples were automatically separated by index.

Subsequently, the resulting reads were mapped to the reference sequence (locus JAPRAS010000022.1:11775–12839) using the minimap2 algorithm.^37^ The quality and structure of the resulting alignment were verified through visualization in IGV.^38^ Finally, the consensus sequences were generated using the samtools toolkit.^39^

### 2.7 Databases and software

The original amino acid sequence of cytochrome b (CYTb) from *Tetranychus urticae* was obtained from GenBank (ID: WQH82754.1). A search for homologous structures was performed using BLASTp with default parameters (word size=5, BLOSUM62 matrix, gap opening/extension penalty of 11/1) against the Protein Data Bank (PDB). Chicken complex III (PDB ID: 1BCC) was used as a reference. The new structure of CYTb from TSSM was predicted using AlphaFold2 with default parameters, using 1BCC as the template. Molecular visualization and structural analysis were conducted in PyMOL 3.0.0.^40^ For modeling subunit complexes, MODELLER 10.6 (Conda version) was used.^41^

### 2.8 Methods of bioinformatics analysis

#### Homologous Modeling

Using BLASTp, homologues of the cytochrome c1 subunit (XM_015925946) and heme protein (XP_025016154.1) were identified in *T. urticae*. The sequences were aligned to the corresponding structures from 1BCC, with unnecessary N-terminal fragments removed. For mutation analysis, structural variants were generated in three ways: (1) all mutations combined, (2) each mutation separately, and (3) pairs of linked mutations.

#### Search for Coevolving Residues

Using BLASTn (default parameters), sequences were searched against a database of mites from the same genus. The sequences listed in the table and the sequence WQH82754.1 served as the initial queries for the search. Deduplicated results and sequences with known mutation effects were then included in the analysis.

Correlating positions and binding pockets were identified using visual CMAT with default parameters, using the AlphaFold2 prediction as the input structure.^42^

#### Determination of Membrane Topology

The positions of transmembrane domains were predicted using DeepTMHMM 1.0.42 based on the sequence.^43^ These predictions were then refined using OPM 3.0 (AWS version), analyzing both with and without membrane curvature.

#### Validation of Results

The quality of the models was assessed by comparing them with the experimental structure (1BCC) and conducting steric clash analysis in PyMOL.

For mutant variants, we checked for preservation of the overall fold and the absence of steric overlaps, except in the case of G126S, where the mutation was found to be structurally incompatible.

#### Computational Resources

Structure prediction was performed using the Google Colab AlphaFold2 interface.^44^

## 3 Results

### 3.1 Structural modeling of cytochrome b from *Tetranychus urticae*

Since the alleles associated with resistance to bifenazate in Russian two-spotted spider mite populations have not been studied, we first compiled a list of mutations documented in the literature that confer resistance to this substance. We then modeled the protein structures to identify which mutations might significantly impact their interaction with the active ingredient.

The structure of *T. urticae* cytochrome b (CYTb) was predicted using AlphaFold2 and exhibits high similarity to chicken cytochrome b (PDB ID: 1BCC; RMSD = 1.2 Å).

Two subunits were identified in complex with cytb:

- Cytochrome c1 (homologue: XM_015925946), which was successfully modeled based on the chicken template.
- Heme protein (homologue: XP_025016154.1), whose structure was predicted with lower confidence (pLDDT < 70 for the N-terminal domain).

### 3.2 Analysis of mutations of *cytb* at the protein level

The desired mutations were introduced into the structure using PyMod (Fig. 1(a)). The G126S substitution created a steric conflict that prevented its stable integration into the structure. The mutations do not interact with neighboring subunits, as the minimum distance to cytochrome c1/heme protein is greater than 8 Å.

**Fig. 1.**
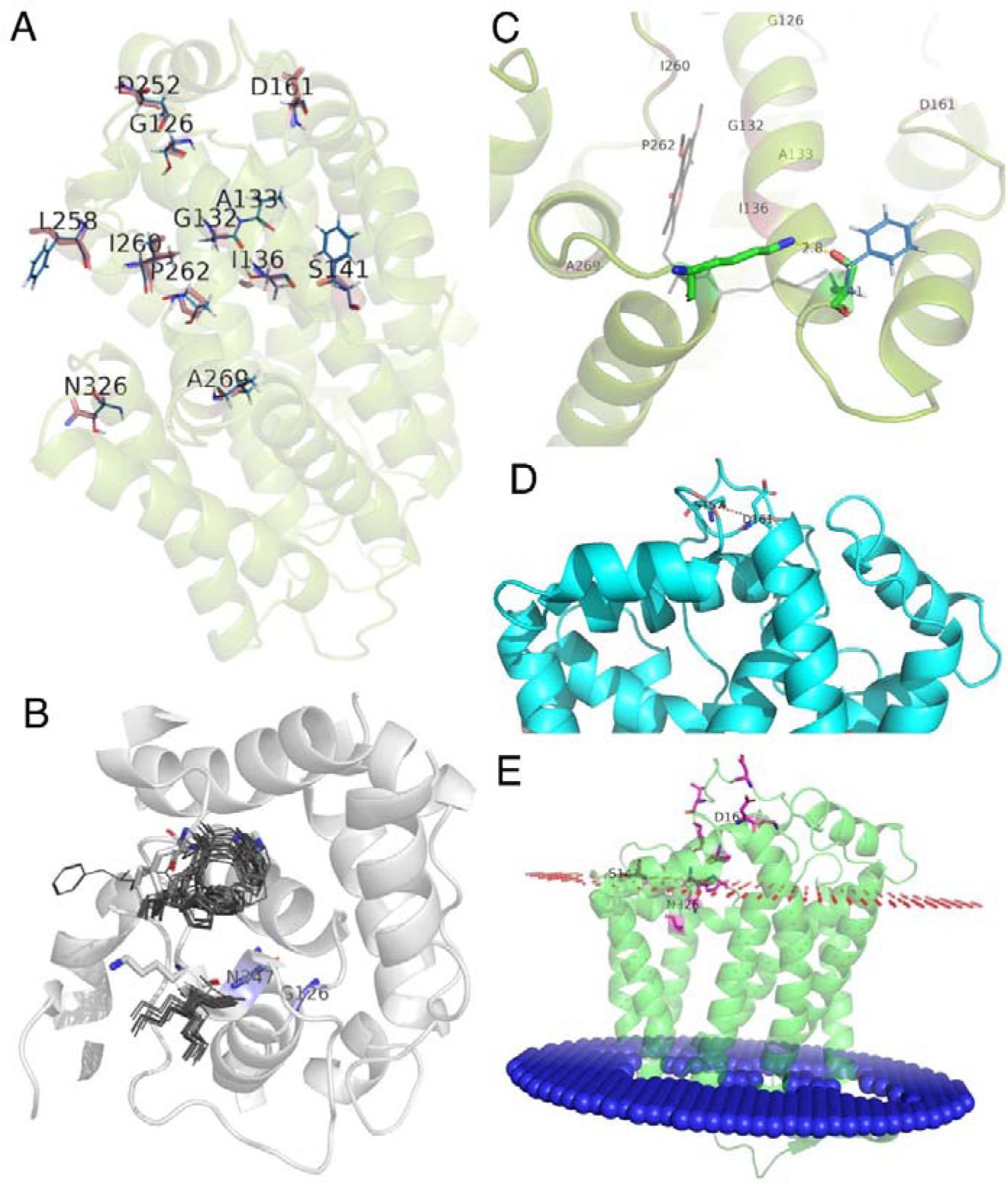
Spatial location of resistance-associated mutations in cytochrome. **b.** (a) Location of candidate residues relative to cytochrome b. (b) Predicted conformations of selected residues in 20 G126S mutants. Wild-type conformations are shown as sticks. Straightforward mutation G126S in original structure leads to a clash with N247. (c) Interaction between S141 and K279. This interaction is disrupted by the S141F mutation; however, cation–π interaction between K279 and F141 may still occur. (d) Coevolving residue pair D161–S157. (e) Position of mutations relative to membrane topology. Unsurprisingly, mutations are found on the positive side of the membrane. N326 and A269 are located inside the membrane. S141 is predicted to lie close to the phosphodiester layer; S141F mutation is a hydrophilicity-hydrophobicity switching.

Additionally, the structures of the mutant proteins were predicted using AlphaFold. The mutant variants—single, paired, and full set—did not exhibit significant structural rearrangements, with an average RMSD of less than 0.8 Å relative to the native model (Supplementary File S1, S2).

Twenty AlphaFold structures were obtained for the mutant protein (4 seeds, max_msa 64:128). Residues within 15 Å of residue 126 were tested, excluding residues 126 and 247. Residues 239–242 and 245 changed their conformation in all mutated structures, exhibiting at least two different orientation types. These amino acids do not appear to make contact with the ligand, heme, or other subunits and seem to comprise a loop, predicted with a low level of certainty (Fig. 1(b)).

Approximately 20 additional residues differ in states between the various structures; however, none of these were found to be significant in any other context.

### 3.3 Coevolution and membrane topology

Coevolutionary Analysis. VisualCMAT revealed significant evolutionary correlations among 345 residues (Supplementary Table S3). Notably, only the 161st residue, which correlates with the 157th, has been previously identified as playing a role in resistance (Fig. 1(d)). VisualCMAT also did not predict the locations of resistance-associated mutations within the binding pockets (Supplementary File S4).

Membrane Topology. Deep TMHMM and OPM predicted eight transmembrane helices, which is in agreement with previous findings (Fig. 1(e)).

The curvature of the membrane did not influence the positioning of mutations, as all are situated on the positive side of the membrane. The substitution of hydrophilic serine with hydrophobic phenylalanine near the predicted membrane surface may lead to slight changes in complex conformation or positioning (Supplementary File S5).

A summary of the protein structure modeling results is provided in Table 2.

**Table 2.**
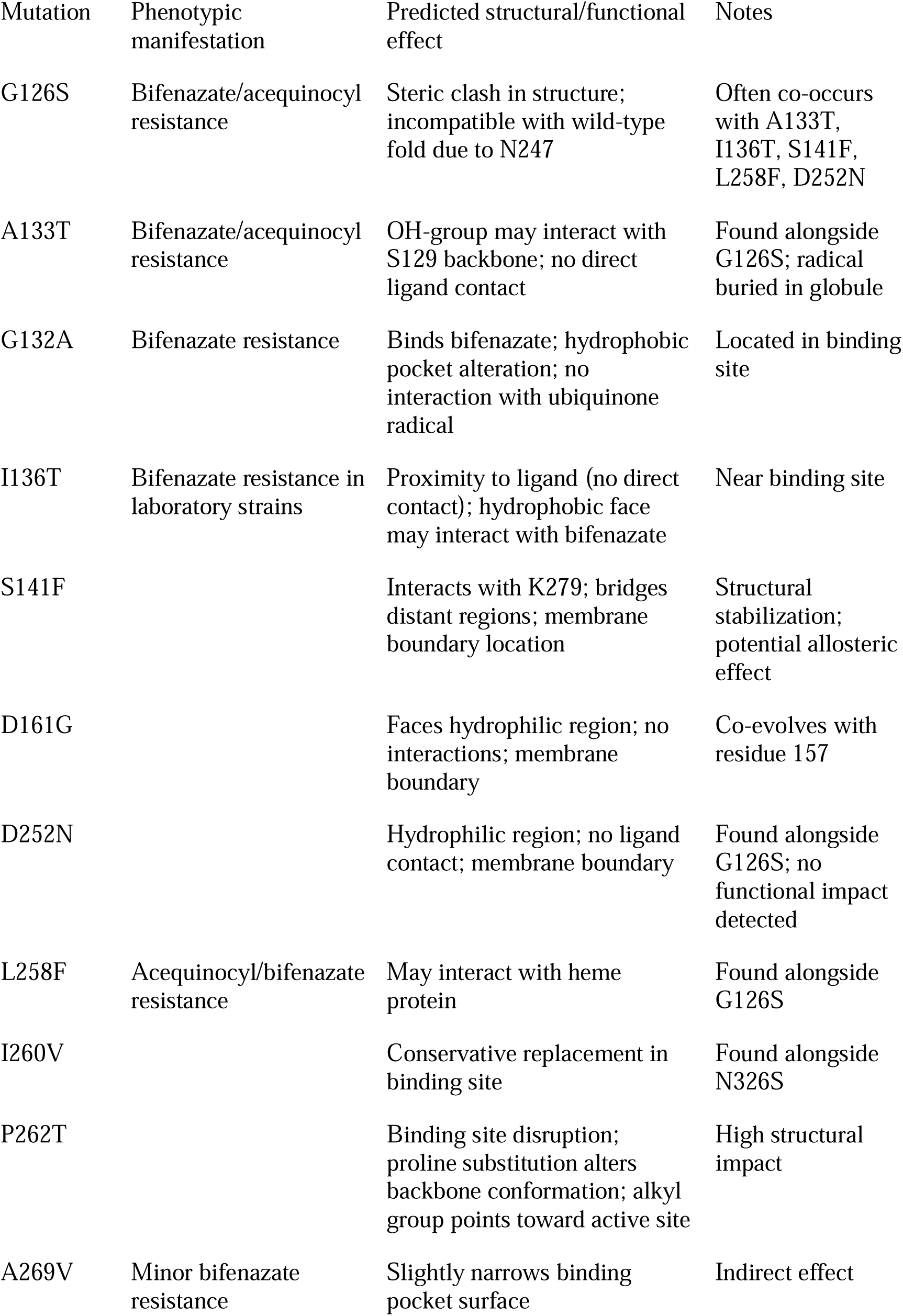
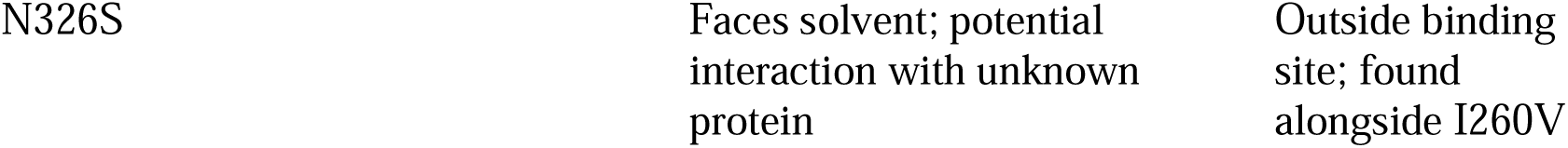
Structural changes in cytb gene mutants in two-spotted spider mite (*Tetranychus urticae* Koch)

Our data align with the molecular docking results obtained by Kewedar and co-authors,^45^ which indicated that the G132A mutation alters the structure of the binding pocket.

Thus:

Mutation-induced structural changes in cytochrome b do not account for stability through direct effects on protein-protein interactions. Instead, they suggest conformational changes within the protein itself (G126S) or in the ligand-binding pocket (G132A, I260V, P262T).

The identified coevolutionary patterns and membrane localization provide new targets for functional studies, with amino acid residues 161 and 157 being particularly noteworthy.

### 3.4 Search for the presence of SNP G126S in the mitochondrial genomes of mites from the resistant culture of VIZR

To detect the G126S mutation in the CYTB gene, we employed a diagnostic system previously described in the literature,^36^ which included both allele-specific primers and primers that amplify a gene region regardless of the mutation’s presence. Analysis of 32 *T. urticae* females (19 from the resistant line (R line) and 13 from the susceptible line (S line)) revealed 17 individuals carrying the G126S mutation. The threshold cycle (Ct) values for both primer types allowed us to estimate the relative copy number of the mutant alleles. Given that the mitochondrial genome exists in multiple copies (ranging from hundreds to thousands of copies per cell),^46^ significant variability in the frequency of mutant alleles was anticipated. In fact, the average difference in threshold cycles (ΔCt) between reactions with allele-specific and universal primers was 9.4 cycles, which, assuming equal reaction efficiency, corresponds to a proportion of the mutant allele of approximately 0.15%. The low frequency of mutant alleles was likely associated with an insufficient concentration of the selective agent used.

### 3.5 Estimation of mutant allele frequency using Oxford Nanopore sequencing

To estimate the allele frequency of the single-nucleotide substitution corresponding to the G126S mutation (G>A substitution at position 376), sequencing was conducted on a diagnostic fragment of the CYTB gene, amplified using the primers BifUnM-F and BifUnM-R. The analysis showed that the frequency of the allele with the G126S mutation was less than 1 percent (226 out of 35,895 reads). No other resistance-inducing substitutions described in the literature were identified, nor were any associated substitutions found during the coevolutionary analysis. In the analyzed region of the gene, three previously undescribed synonymous substitutions were identified that do not alter the amino acid sequence of the protein (441A>G, c.465G>A, and c.624T>C – numbering from the start of the coding sequence) (Supplementary File S6). These substitutions can be used for molecular labeling of the laboratory culture of spider mites from the VIZR collection.

### 3.6 Selection at elevated concentrations of bifenazate

To assess the role of the identified allele with the G126S substitution in the development of bifenazate resistance, we conducted selection treatments on beans infected with individuals from the R line over the course of a year. The concentration of the drug was gradually increased throughout the series of treatments. Upon completion of the experiment, we performed toxicological testing on the crop subjected to the selection treatments using both the diagnostic and maximum concentrations of bifenazate. At the diagnostic concentration, the average mortality rate was 18.4%, indicating the development of resistance to the drug. At a concentration of 0.031%, survival was limited to just a few individuals.

### 3.7 Estimation of mutant allele frequency after selection at elevated concentrations of bifenazate

After a series of selection treatments with stepwise increasing drug concentrations, real-time PCR was repeated using the diagnostic system described above.^36^ The difference in threshold cycles between reactions with allele-specific and universal primers (ΔCt) decreased to 1.75, indicating an increase in the proportion of the resistant allele.

Similar to the previous analysis, Oxford Nanopore sequencing was performed on laboratory mite cultures after completing the bifenazate treatments. Before selection (initial culture), the wild-type allele dominated at the analyzed position, accounting for 99% of the total 35,895 reads. After a series of selection treatments with bifenazate, the allele ratio reversed. Out of 8,056 reads, the mutant allele was detected in 90% (7,272), while the wild-type allele was present in 10% (770) (Supplementary File S7). Thus, the selection treatments resulted in a significant increase in the proportion of the mutant allele, establishing its dominance (Fig. 2). No other mutations conferring resistance were identified. The data indicate rapid fixation of the resistant allele under chemical selective pressure.

**Fig. 2.**
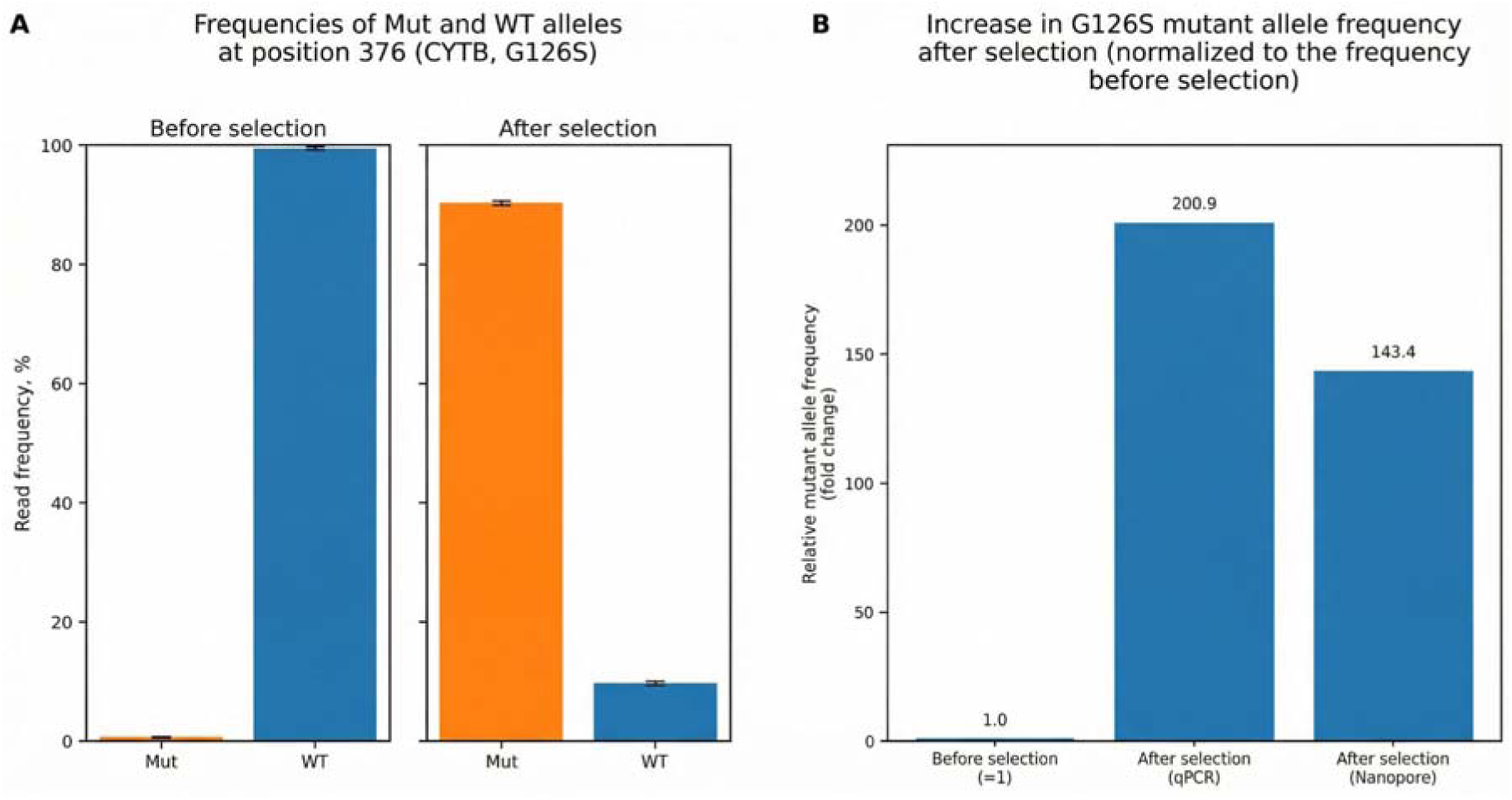
Allele frequencies of the *cytb* gene before and after bifenazate selection. (a) Allele frequencies estimated based on Oxford Nanopore sequencing results. (b) Comparison of mutant allele frequency estimates obtained by qPCR and nanopore sequencing.

## 4 Discussion

This study demonstrates, for the first time, the presence of the G126S mutation in a laboratory culture derived from the Russian TSSM population. The frequency of the allele containing this mutation increased by two orders of magnitude under strong selective pressure from bifenazate.

This assessment was made using two methods: Oxford Nanopore sequencing and real-time PCR. No other mutations leading to amino acid substitutions associated with bifenazate resistance were detected in the diagnostic gene fragment, which supports the role of G126S mutation in mediating resistance to the substance in question. This conclusion is further bolstered by experiments modeling the protein structures of both the normal and mutant alleles. Our findings align with the data presented by Shi et al. ^47^ and contradict the views of some other authors. ^34,35^ This discrepancy may stem from the different genetic backgrounds in which mutations manifest in TSSM from various populations. The *cytb* gene product is a protein that functions as part of a complex multisubunit assembly. Consequently, we can hypothesize the existence of compensatory mutations outside the gene under investigation that could affect its functionality. To elucidate the mechanisms of the mutation’s action, more detailed studies are needed that extend beyond the structure of the gene and cytochrome b protein.

Furthermore, it is not only important to detect the G126S substitution in the mite population but also to assess the frequency of the mutant allele relative to the wild-type variant. Our experiment showed a two-order-of-magnitude increase in the frequency of the mutant allele in the culture maintained at high concentrations of the active ingredient.

## 5 Conclusions

This study provides the first molecular genetic characterization of bifenazate resistance in a Russian population of the two-spotted spider mite, *Tetranychus urticae*. We successfully established resistant and susceptible isogenic lines and applied structural bioinformatics, targeted genotyping, and next-generation sequencing to elucidate the resistance mechanism. Our key finding is the identification and validation of the G126S mutation in the mitochondrial cytochrome b gene as a central factor conferring resistance in this specific genetic background. Structural modeling indicated that this substitution causes significant steric conflict, likely destabilizing the protein fold or its interaction with bifenazate.

Critically, longitudinal selection experiments with escalating bifenazate concentrations demonstrated a direct correlation between selection pressure and the frequency of the G126S allele, which increased from less than 1 % to 90 % dominance in the population. This rapid fixation under chemical stress strongly supports a causal role for G126S in the observed resistance phenotype. Furthermore, the absence of other known resistance-associated mutations in the analyzed *cytb* fragment suggests that, contrary to some reports from other global populations, G126S alone can be a primary determinant of resistance in certain genetic contexts.

Our findings reconcile conflicting literature by emphasizing the population-specific nature of resistance evolution. The discrepancy regarding the indispensability of G126S likely stems from differing genetic backgrounds, where its effect may be modulated by unlinked compensatory mutations or synergistic metabolic mechanisms not examined here. This research underscores the necessity of monitoring local pest populations for resistance alleles and confirms the G126S mutation as a potent and selectable marker for bifenazate resistance. Future work should focus on comprehensive genomic and metabolomic analyses to identify potential contributing factors outside the *cytb* gene and to develop robust diagnostic tools for resistance management in agricultural settings.

## Acknowledgements

This work was performed using the equipment of the Research Park of St. Petersburg University.

